# VISION — an open-source software for automated multi-dimensional image analysis of cellular biophysics

**DOI:** 10.1101/2024.03.29.587344

**Authors:** Florian Weber, Sofiia Iskrak, Franziska Ragaller, Jan Schlegel, Birgit Plochberger, Erdinc Sezgin, Luca A. Andronico

## Abstract

Environment-sensitive probes are frequently used in spectral/multi-channel microscopy to study alterations in cell homeostasis. However, the few open-source packages available for processing of spectral images are limited in scope. Here, we present VISION, a stand-alone software based on Phyton for spectral analysis with improved applicability. In addition to classical intensity-based analysis, our software can batch-process multidimensional images with an advanced single-cell segmentation capability and apply user-defined mathematical operations on spectra to calculate biophysical and metabolic parameters of single cells. VISION allows for 3D and temporal mapping of properties such as membrane fluidity and mitochondrial potential. We demonstrate the broad applicability of VISION by applying it to study the effect of various drugs on cellular biophysical properties; the correlation between membrane fluidity and mitochondrial potential; protein distribution in cell-cell contacts; and properties of nanodomains in cell-derived vesicles. Together with the code, we provide a graphical user interface for facile adoption. We anticipate that VISION will find a broad range of applications in different fields of biology, spanning from molecular and tissue biology to immunology and biophysics.

**Summary statement:** VISION, an open-source software, enables high throughput and correlative analysis of cellular biophysical properties.

## Introduction

Biophysics aims at understanding the correlations between physical changes in cell and biological processes (Horwitz, 2016). For example, researchers have demonstrated how remodelling of membrane biophysical properties, including polarity, tension, fluidity, and depolarization, can influence processes such as cell proliferation and migration (Atilla-Gokcumen *et al*., 2014; Braig *et al*., 2015; Edmond *et al*., 2015; Kunduri, Acharya and Acharya, 2022) and immune response (Rentero *et al*., 2008; Rudd-Schmidt *et al*., 2019; Tello-Lafoz *et al*., 2021). Recent advances in technology for high-throughput characterization of cell biophysics, have highlighted the importance of such biophysical remodelling also in diseases (Andronico *et al*., 2024). Among others, environment-sensitive probes are often used to investigate alterations in cellular biophysics. These probes undergo changes in their photophysical properties (e.g., emission intensity and/or spectrum, fluorescence lifetime) in response to changes in the surrounding lipid composition or bilayer hydration (García-Calvo *et al*., 2022; Ragaller *et al*., 2022, 2023). Typical examples of membrane-intercalating dyes, which undergo an emission red-shifting upon increase in membrane fluidity, are Laurdan, NR12A/S and Pro12A (Amaro *et al*., 2017; Danylchuk *et al*., 2020; Niko and Klymchenko, 2021). These probes can be chemically functionalized to target either the plasma membrane (PM) or the lipid bilayer of different subcellular compartments (Klymchenko, 2023). Thus, fluorescence microscopy is often used to study 2D- and 3D-dynamic remodelling of cell biophysics, specifically under spectral or multi-channel modality. Indeed, by acquiring a greater extent of the emission spectrum, the resolution and sensitivity of measurements can be enhanced (Sezgin *et al*., 2015). Then, the image analysis usually requires some form of mathematical operations to derive a practical redout of the biophysical properties under investigation. This could be as simple as calculating the ratio between two intensities (*e*.*g*., to measure mitochondrial membrane depolarization *via* JC-1 dye) (Sivandzade, Bhalerao and Cucullo, 2019), or it might involve a more elaborated equation/fitting, for instance as for the calculation of the generalized polarization (GP) which is used to determine membrane fluidity (Yu *et al*., 1996). Several open-source imaging programs have been developed to analyse object properties such as morphology, overall fluorescence intensity, number of particles *etc*. These software are often applicable either to certain geometry such as spherical vesicles (Stirling *et al*., 2021; Buren, Koenderink and Martinez-Torres, 2022) or specialized in analysing specific subcellular compartments, such as the PM or the cell nucleus (Barry *et al*., 2022). The options decrease even further when one needs to analyse cellular biophysical properties which requires mathematical operations. Here, we introduce a new tool for analysis of 2D/3D multi-channel/spectral images, implemented in Python and featuring a user-friendly graphical interface (GUI). In addition to canonical intensity-based analysis, our software enables for profiling of several cell biophysical parameters (*i*.*e*., membrane fluidity, mitochondria membrane potential *etc*.) in an automated fashion and from multiple image formats. The possibility to calculate multiple biophysical readouts is achieved through the ability to perform customizable mathematical operations on the different acquired channels. Some of the key features are independent masking of PM and cytosol, whole-image *vs* single-object analysis, morphological characterization, high-resolution membrane profiling as well as cell linearization. We demonstrate the advantages of our software in four different biological scenarios: *i)* characterization of phase stability in cell-derived vesicles; *ii)* correlation between mitochondria health and fluidity of the plasma membrane under pathological settings; *iii)* mapping of the protein accumulation and clustering in reconstituted immune synapses and *iv)* profiling of the membrane fluidity in phase separated vesicles at submicron resolution. Therefore, we believe our software represents a complementary tool for researchers who are interested in investigating spatiotemporal aspects of biophysical and metabolic remodelling caused by cellular processes and external stimuli.

## Results

### Software overview

Most of the functions utilized in our software sourced from common open-source packages (*i*.*e*., scipy, scikit-learn, matplotlib), whereas others were developed by us to perform specific tasks during the analysis. VISION is highly versatile in terms of input image types which can be analysed. It supports various file formats (Fig. 1) and it can handle multi-dimensional images, including t-stack, z-stack, and t-z-stack formats. Additionally, VISION offers batch processing capabilities. The software is designed to take advantage of the enhanced detail in measuring biophysical properties in cells which derives from working with spectral images (Aron *et al*., 2017). Therefore, it can process multi-dimensional images with one and up to *n* λ-channels. A key aspect of analysing microscopy images is the creation of a proper mask. In this regard, VISION offers users significant flexibility, enabling independent masking of the plasma membrane and cytosolic parts, as well as performing mask morphological reconstruction after thresholding. After the mask has been created, the user can choose which biophysical parameter (which we refer to as β-value) to calculate (*via* a customized equation), and whether to perform the analysis on the image as a whole or on individual objects. Thus, several outputs are generated such as *i)* the reconstructed emission spectrum, *ii)* the β-value colour-coded image, *iii)* a histogram of the β-value distribution and *iv)* the phasor plot of β-value distribution. The single-object mode allows for the extraction of additional descriptive statistical and morphological information on individual objects. It also enables the export of a spectrally reconstructed image of either the individual object’s membrane/cytosol or the linearized object. Finally, all data sets are saved and exported into an Excel file to enable further downstream analysis. With its numerous implemented features, our software can be utilized for quantitative analysis of a variety of different samples such as single particles, vesicles or cells.

**Fig. 1.**
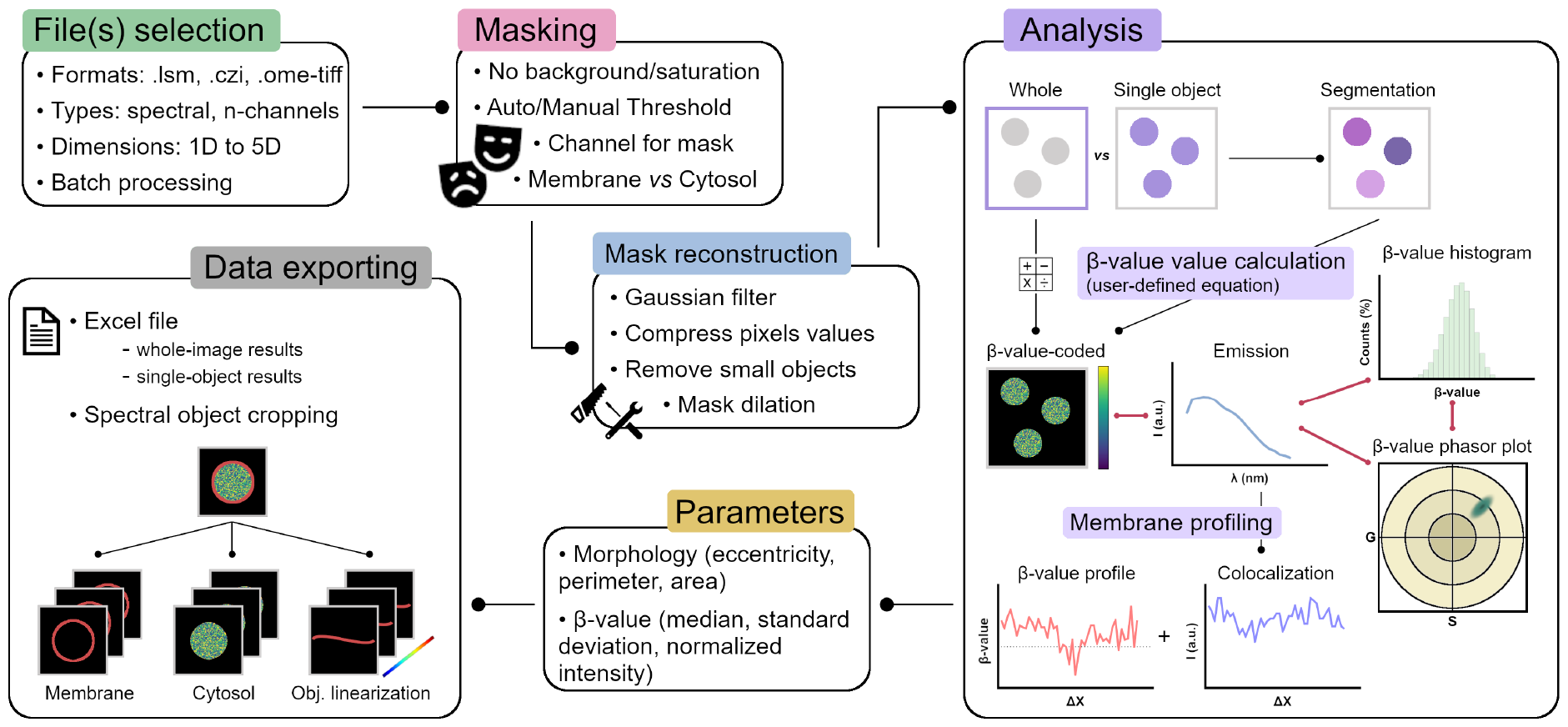
Illustration of the VISION general workflow. The different boxes show the main features of the VISION software.

### Masking

In assessing the biophysical properties, the incorporation of noisy pixels (*i*.*e*., background fluctuations) in the calculation of the β-value could potentially have pronounced impact on the measurement accuracy. These pixels introduce uncertainty which propagates and increases due to the mathematical operations typically performed to derive the β-value. For this reason, the software offers the capability to specify a particular signal-to-noise ratio for filtering out background pixels before image thresholding, and it autonomously eliminates saturated pixels. The analysis of individual objects by VISION relies on creating a skeleton from the thresholded image. Thus, for membranes, it is fundamental to obtain a mask where the membrane is a continuum of pixels. This can be achieved by performing different implemented operations on the mask. For instance, the inserts in Fig. 2A, compare the results obtained when *i)* applying only an automatic thresholding *via* Otsu algorithm, *ii)* applying a Gaussian filter before the automatic thresholding and *iii)* compress the pixels values in between the Gaussian filter and the thresholding *via* Otsu. Specifically, the use of our custom-defined function for pixels compression (Fig. S1) allows to threshold areas with different brightness *via* Otsu algorithm, thus preserving the automation of thresholding during batch processing. This approach greatly simplified the masking procedure compared to other software where multiple masks are required to deal with images with heterogenous brightness (Stirling *et al*., 2021). The user can also perform morphological reconstruction of the mask to remove small clusters before the segmentation step (insert 4) and decide the channel (for spectral images) to use for image thresholding. Finally, within the GUI, users have the option to define either global or slice-specific parameters (*e*.*g*., for t-stacks, *etc*.) for masking before initiating batch analysis.

**Fig. 2.**
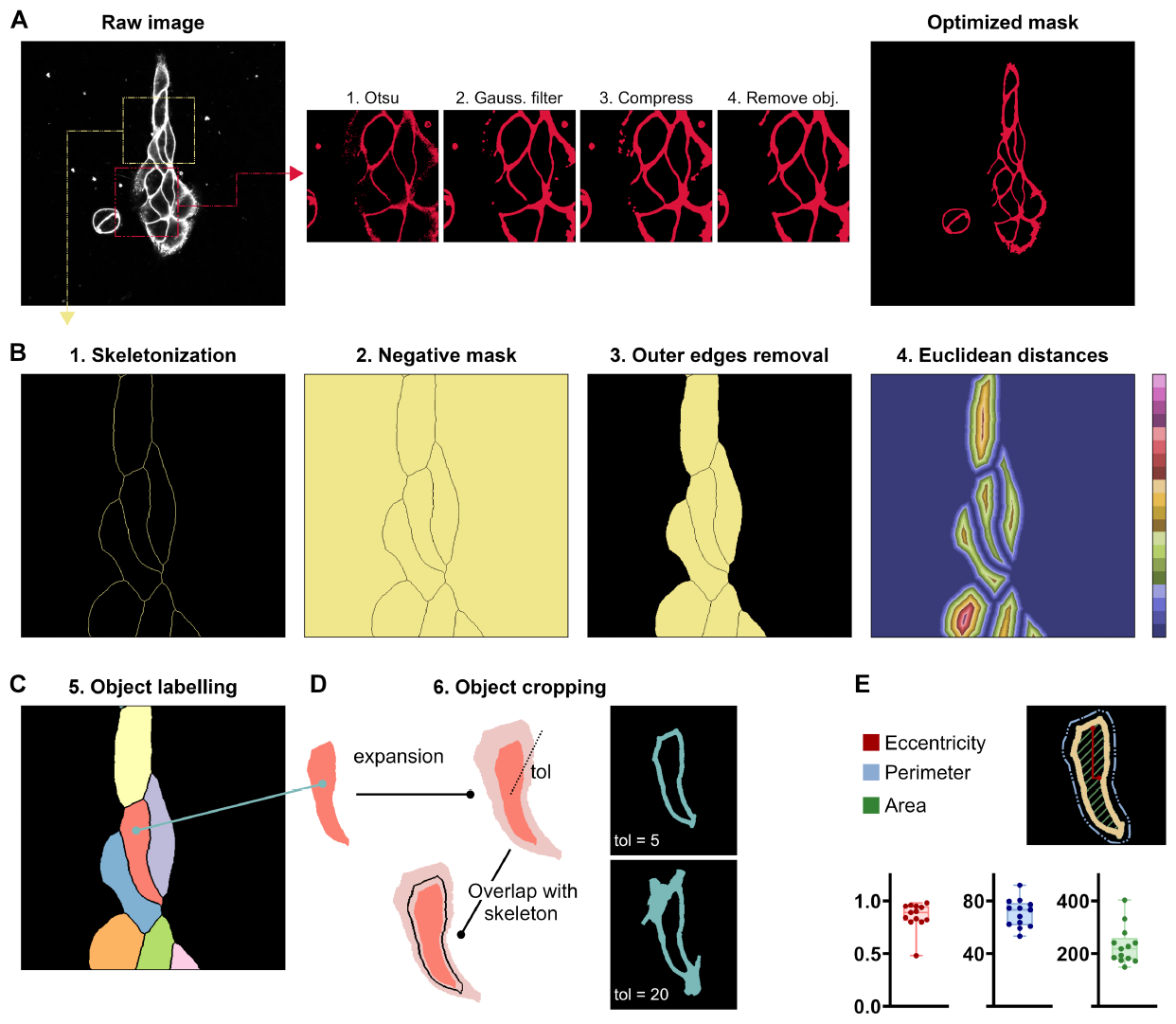
Working principles of image segmentation. (A) Optimization of the mask for the analysis of a parameter on cells. Starting from the raw image, different independent operations can be performed on the preliminary mask for optimization. (B-C) Working principle for the image segmentation. Steps 1 to 5 show the operations performed on the optimized mask to obtain a final image of individually labelled objects. (D) Consecutive steps for the cropping of individual objects. Depending on the tolerance parameter (tol) the user can expand/reduce the cropped object. (E) Object-specific morphological parameters calculated by the software.

### β-value calculation

Once the mask has been optimized, the software provides the option to select up to four channels for calculating the parameter (β-value) of interest. Additionally, users can define a mathematical equation for determining the β-value. This is independent for the membrane and the cytosol calculation, thus allowing for simultaneous characterization of multiple biophysical/metabolic properties and correlative analysis. As a result, it generates a reconstructed pseudo-coloured image displaying the pixel-wise β-values for membrane and cytosol parts individually. The equation supports all the main algebraic operations and pixels which return erroneous values are automatically removed from the analysis. Statistical descriptors for the β-values (*i*.*e*., median, standard deviation) are generated from either the whole image or individual objects, depending on the mode of operation.

### Segmentation

Single-object analysis from microscopy images usually requires two distinct steps: the image segmentation (especially when working with clustered objects such as adherent cells) followed by the object identification/labelling. Available software recur to either morphology-related (Buren, Koenderink and Martinez-Torres, 2022) or AI-based algorithms (Berg *et al*., 2019; Körber, 2022) to perform object detection. In VISION, object detection is accomplished through a six-step algorithmic process (Fig. 2B-C). This process can be applied to images containing a mixture of isolated and clustered objects, with shapes ranging from simpler to highly irregular. First, a raw skeleton is derived from the optimized mask and subjected to a customized function for removal of side branches; the skeletonized mask is converted into a negative mask where the inner part of the object is filled, and the outer edges are removed. This operation ensures that objects sitting at the image edges (*i*.*e*., whose membrane is cropped) will be ignored during the analysis. The mask of filled objects is then converted into the corresponding mask of pixel-to-pixel Euclidean distances. At this stage, a user-defined tolerance value (*tol*_*0*_) will be used to threshold only those distances above *tol*_*0*_, thereby causing a grater or smaller retraction from the vesicle’s skeleton. Subsequently, shrunken vesicles will be progressively labelled after undergoing pixel clustering via the DBSCAN algorithm. DBSCAN is a density-based algorithm that performs effectively on arbitrarily shaped vesicles and without the need to predefine the number of clusters (Martin Ester, 1996). The last step of image segmentation represents the cropping of individual vesicles *via* consecutive expansion of the shrunk vesicles (according to a second user-define tolerance value *tol*) and overlap with the general skeleton. The software returns a set of morphological parameters for each individual vesicle, such as vesicle’s eccentricity, perimeter and area (Fig. 2D) which are calculated using modules from the open-source scikit-learn package (Pedregosa *et al*., 2011).

### Plasma Membrane profiling

A key feature of the VISION software is the possibility to profile the membrane and track fluctuations of the β-value and/or channel intensities at a resolution which is limited only by the pixel size. This characteristic is useful, for instance, when studying the dynamic rearrangement of biophysical properties at the cell-to-cell contact, to monitor protein accumulation in phase-separated lipid bilayers or protein clustering at the membrane sub-compartments (Lamerton *et al*., 2021). Fig. 3 shows the principles behind the module for membrane profiling applied to two different scenarios: suspension cells (HL-60, Fig. 3A-B) and phase-separated giant unilamellar vesicles (GUVs), both stained with an environment-sensitive probe for membrane fluidity. Our approach relies on creating a 1-pixel wide guiding line centred onto the membrane which will be used to move a 2D integrating element unidirectionally along the bilayer for signal integration. Therefore, it is important to avoid any cytosol contribution when analysing cells (Fig. 3B), or to ensure continuity of the masked membrane when analysing phase-separated synthetic vesicles with areas of different brightness (Fig. 3E) during the masking step. Once the mask is created, it will undergo skeletonization, morphological reconstruction of the skeleton and unidirectional re-ordering of the line’s pixels coordinates, *via* a customized function (Fig. 3C). In case of clustered vesicles, the software will automatically detect the intersection nodes and identify the different segments which constitute the clustered vesicles’ or cells’ membrane. Finally, a moving average filter, shaped according to user-defined parameters, will traverse along the ordered skeleton (Fig. 3F) and generate trajectory curves indicating the average intensity/β-value. By adjusting the shape and size of the moving average filter, users can obtain smoother trajectories if desired (Fig. 3G), although at the cost of spatial resolution. Common methods for membrane profiling involve calculating the integrated signal from segments of the membrane by slicing it into portions of equal angular amplitude (Buren, Koenderink and Martinez-Torres, 2022). Although this method works for spherical objects, it might yield inaccurate results on cells with irregular shapes. On the other hand, our approach ensures the same integrating area, thus leading to a greater accuracy even when working in the presences of membrane irregularities such as budding, invagination *etc*. The GUI also offers the option to perform a recentring of the guiding line onto the membrane (useful when analysing membranes which present clusters or aggregated proteins) (Fig. S2) and the colocalization between the β-value and the signal from a third channel of interest.

**Fig. 3.**
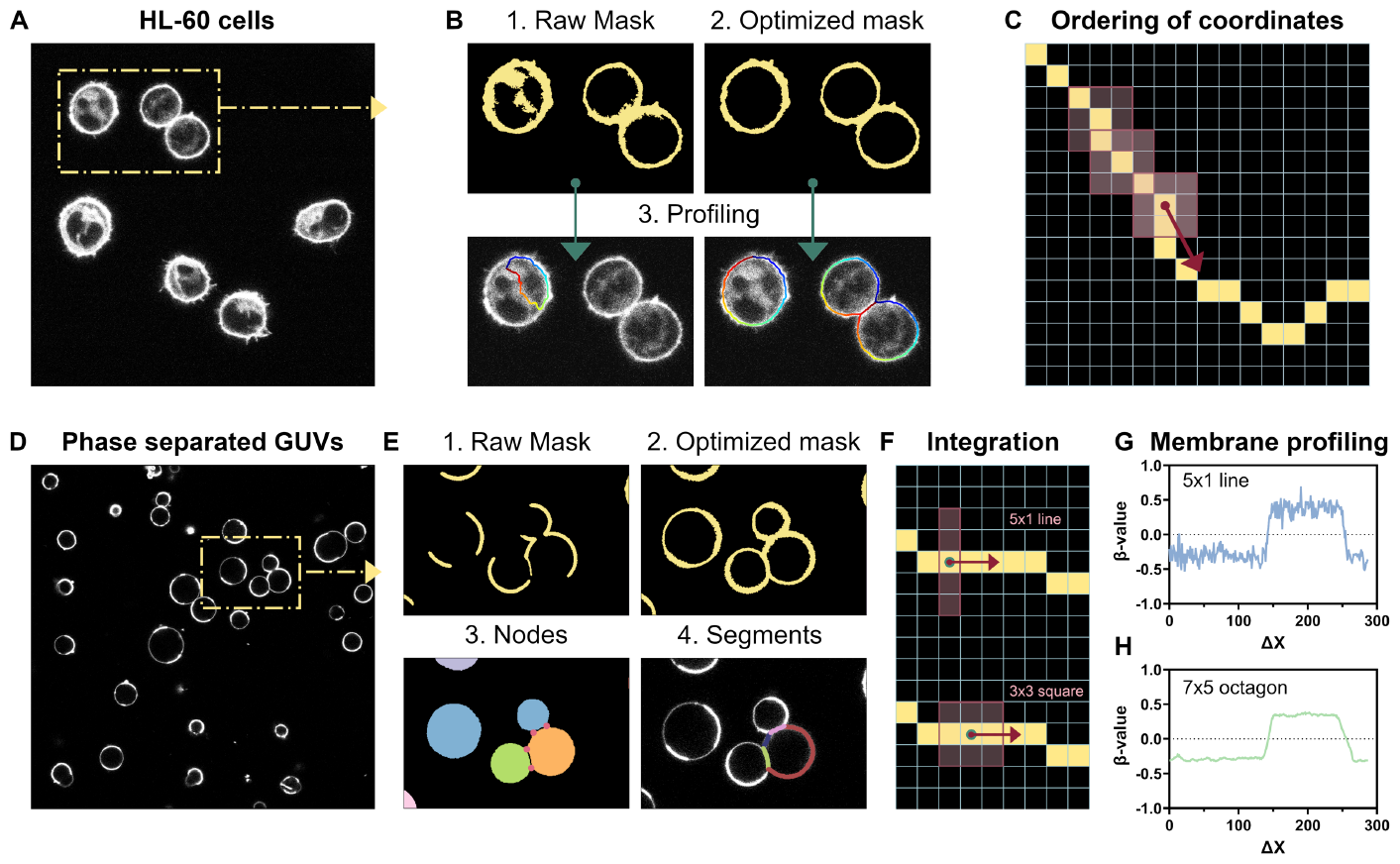
Profiling of membranes. (A) Raw image of Human Leukemia cell lines (HL-60) suspension cells stained with an environment-sensitive probe (Pro12A) used for measurement of membrane fluidity. (B) The effect of mask optimization on the profiling of the plasma membrane. The raw mask is obtained by applying an automatic threshold *via* the Otsu algorithm, whereas the optimized mask is derived by choosing a channel with low cytosolic background for thresholding and applying the function for pixel compression, to obtain a continuous membrane. (C) Schematic for the ordering of the pixel coordinates from the 1-pixel wide line used for the membrane profiling. The line (in yellow) is centred onto the plasma membrane, and a 3x3 pixel square box is slid along the line. (D) Raw image of phase-separated GUVs stained with an environment-sensitive probe (NR12S) used for measurement of membrane fluidity. (E) The effect of mask optimization for membrane profiling and segment detection in clustered vesicles. The raw mask is obtained by applying an automatic threshold *via* the Otsu algorithm, whereas the optimized mask is derived by choosing a channel where the two phases have similar brightness and after applying the function for pixels compression followed by dilation of the binary mask, to obtain a continuous membrane. On the optimized mask, the software detects nodes in clustered vesicles and, if the profiling mode is activated, it profiles individual segments. (F) Schematic for the signal integration during membrane profiling (line vs square). (G-H) The effect of choosing a different integrating element on profiled curves for β-value (*i*.*e*., parameter under investigation). The larger the area of the integrating element the smother the profiled curve. ΔX represents the lateral displacement along the vesicle’s membrane in pixels units.

### Data exporting

For each analysed image, VISION generates an Excel file that is automatically updated after analysis, depending on the specific analysis performed. Column headers are provided to facilitate navigation of the file content. Descriptive statistics on the β-value, the emission spectrum and the β-value histogram are generated from the whole image and, if performing image segmentation, for each individual object detected. Furthermore, when the option for membrane profiling is active, the object-specific trajectories for channels intensities, β-value and additional channel for colocalization are saved, together with the information regarding the number of segments and corresponding length per object (in case of clustered objects).

### Analysis of phase separated vesicles

Thanks to the modules for single object detection and membrane profiling, VISION is very useful when characterizing phase-separated vesicles. In Fig. 4, we present an example of application for analysing phase stability in cell-derived giant plasma membrane vesicles (GPMVs). GPMVs differ from synthetic vesicles, such as GUVs, in that they retain majority of the membrane complexity of their cell precursors (Sezgin, 2022). Thus, we stained GPMVs with Pro12A (which reports on membrane fluidity) and investigated the effect of the chemical deoxycholic acid (DCA) on ordered-disordered phase transition in GPMVs. DCA is a bile acid, naturally found in human body. Its main function is emulsification and solubilization of fats in the body and it has been used as a biological modulating agent in immunology, cancer and cosmetics. It has been shown to stabilize phases with lower membrane order and to predominantly affect the disordered phase fluidity (Zhou *et al*., 2013). As shown in Fig. 4A-C, the addition of DCA had no visible effect on the vesicle morphology, which retained the same rounded shape and size distribution. On the other hand, a shift towards negative values was observed for the generalized polarization (GP) distribution, suggesting that DCA changes the overall membrane fluidity in GPMVs. However, from the histogram of object-specific median GPs (Fig. 4C) it is not possible to conclude whether the observed shift reflects an increase in the total area of disordered domains per vesicle, or an increase in fluidity of the existing domains. To this regard, by profiling the vesicle membrane along the equatorial plane, we were able to trace spatial variation of GP from individual GPMVs and group the GP values measured at each point along the membrane into two distinct sets either above or below a defined threshold (*i*.*e*., the median GP value). Thus, we could observe an increase in the number of pixels (*i*.*e*., membrane areas) with lower membrane fluidity (Fig. 4D) in GPMVs treated with DCA. This evidence supports the hypothesis of a relative increase in the total surface area of low-GP phases compared upon DCA treatment. Furthermore, by plotting values from the two GP sets with and without the bile acid (Fig. 4E), we were able to confirm the results in literature (Zhou *et al*., 2013) which showed that DCA has no effect on the fluidity of ordered phases, but it further decreases the fluidity of the disordered areas. Besides yielding information on the overall extent and GP magnitude of the different phases, profiling *via* VISION can also provide insights upon the morphology and size of domains in vesicles (Fig. 4G-I). Indeed, by looking at the profiled trajectories, it is possible to derive parameters such as, the length of phases, the intra-phase heterogeneity of fluidity (*i*.*e*., fluctuations in GP values) and the gradient of phase separation (from the slopes of GP trajectory). To this regard, we found that the addition of DCA mainly leads to the formation of only two large phases with distinct GP, instead of multiple smaller ones (Fig. 4H-I). For comparison, we also produced GUVs that spontaneously phase separate without the need of additional stabilizers. These synthetic vesicles showed similar morphology to GPMVs (Fig. 4J) but a different efficiency in phase separation. Indeed, as inferable from Fig. 4K, only half of the vesicles showed multiple phases with different membrane fluidity and lengths similar to the case of GPMVs with DCA (Fig. L-M).

**Fig. 4.**
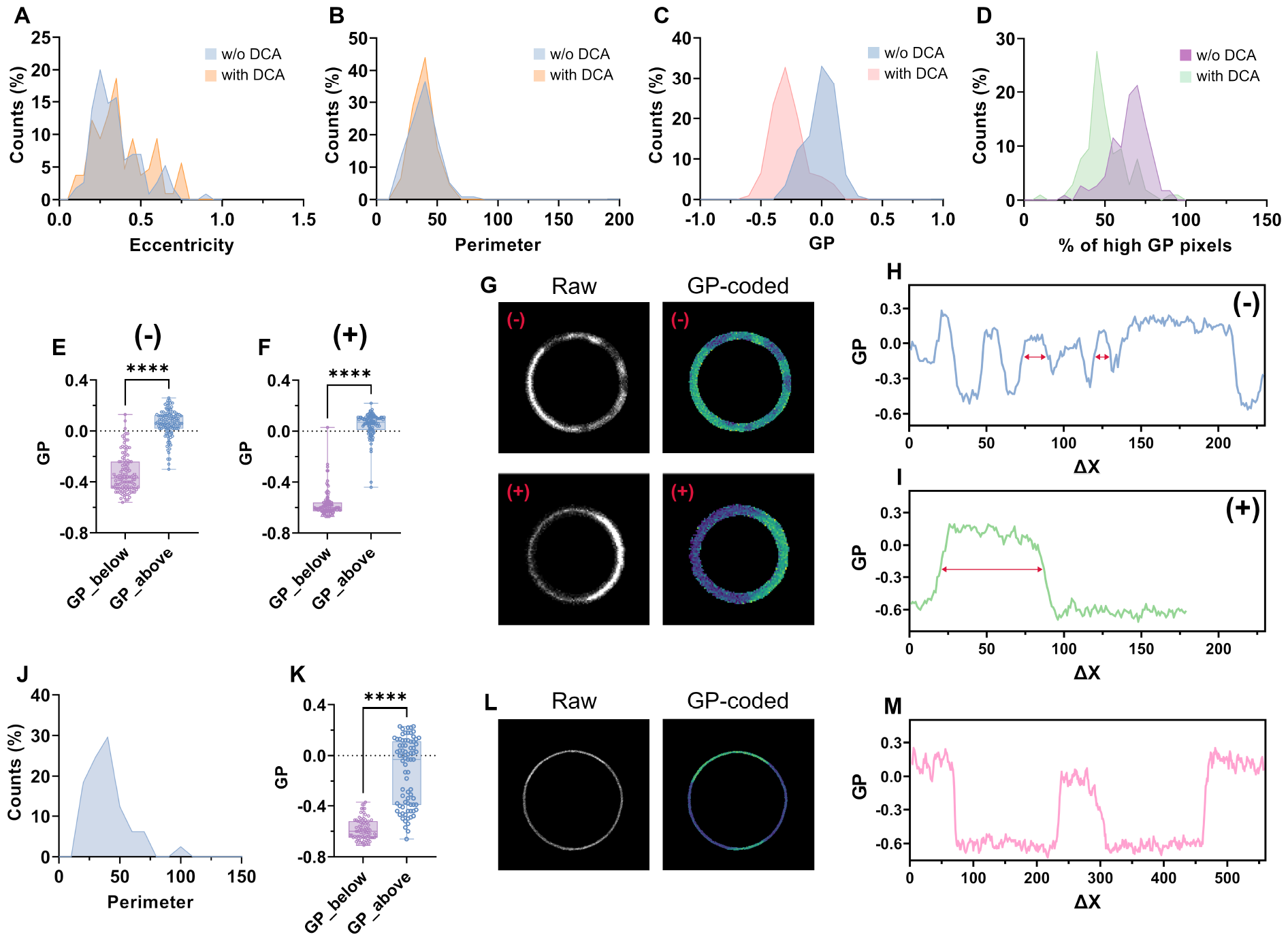
VISION reveals differences in phase stability from synthetic vesicles. (A-D) Distribution of vesicle eccentricity (A), perimeter (B), GP (C) and percentage of high-GP pixels (D) from phase separated GPMVs with and without the addition of DCA. (E-F) Box plots reporting the median GP values of the two subsets of profiled points (falling either above or below the median GP from the profiled trajectory) from individual phase-separated GMPVs (n >100) without (E) or with (F) the addition of DCA. The non-parametric t-test analysis shows significant difference between the two subsets. (G-I) Raw and GP-colour coded representative images and profiled membrane (H, I) of a phase-separated GPMV without (-) and with (+) the addition of DCA. The red arrows highlight differences in phase lengths between the two conditions. (J-K) Distribution of vesicle’s perimeter (J) and box plot (K) reporting the median GP values of the two subsets of profiled points (falling either above or below the median GP from the profiled trajectory) from individual phase-separated GUVs (n > 80). The non-parametric t-test analysis shows significant difference between the two subsets. (L-M), Raw and GP-colour coded representative image and profiled membrane (M) of a phase-separated GUV.

### Correlation of membrane fluidity with metabolic activity

VISION allows for simultaneous and independent characterization of multiple biophysical properties from both the plasma membrane and the cytosol. Thus, we utilized our software to investigate the correlation between remodelling of membrane fluidity and mitochondrial potential under stress. Specifically, we conducted experiments on epithelial-like rat kidney cells (NRK-52E) treated with either β-cyclodextrin or the protonophoric uncoupler carbonyl cyanide m-chlorophenyl hydrazone (CCCP). While cyclodextrin removes cholesterol from the plasma membrane (Zidovetzki and Levitan, 2007), CCCP disrupts the potential of mitochondrial membrane by promoting the opening of the mitochondrial permeability transition pore (Ganote and Armstrong, 2003). As shown in Fig. 5A, we co-stained cells with Pro12A and JC-1 (for mitochondrial membrane potential (Sivandzade, Bhalerao and Cucullo, 2019)) and acquired the intensity from four different channels. The green and red channels were used to detect signal from JC-1 monomers and aggregates (used to mask mitochondria), respectively and calculate the mitochondrial membrane potential as the intensity ratio *I*_*red*_*/I*_*green*_.

**Fig. 5.**
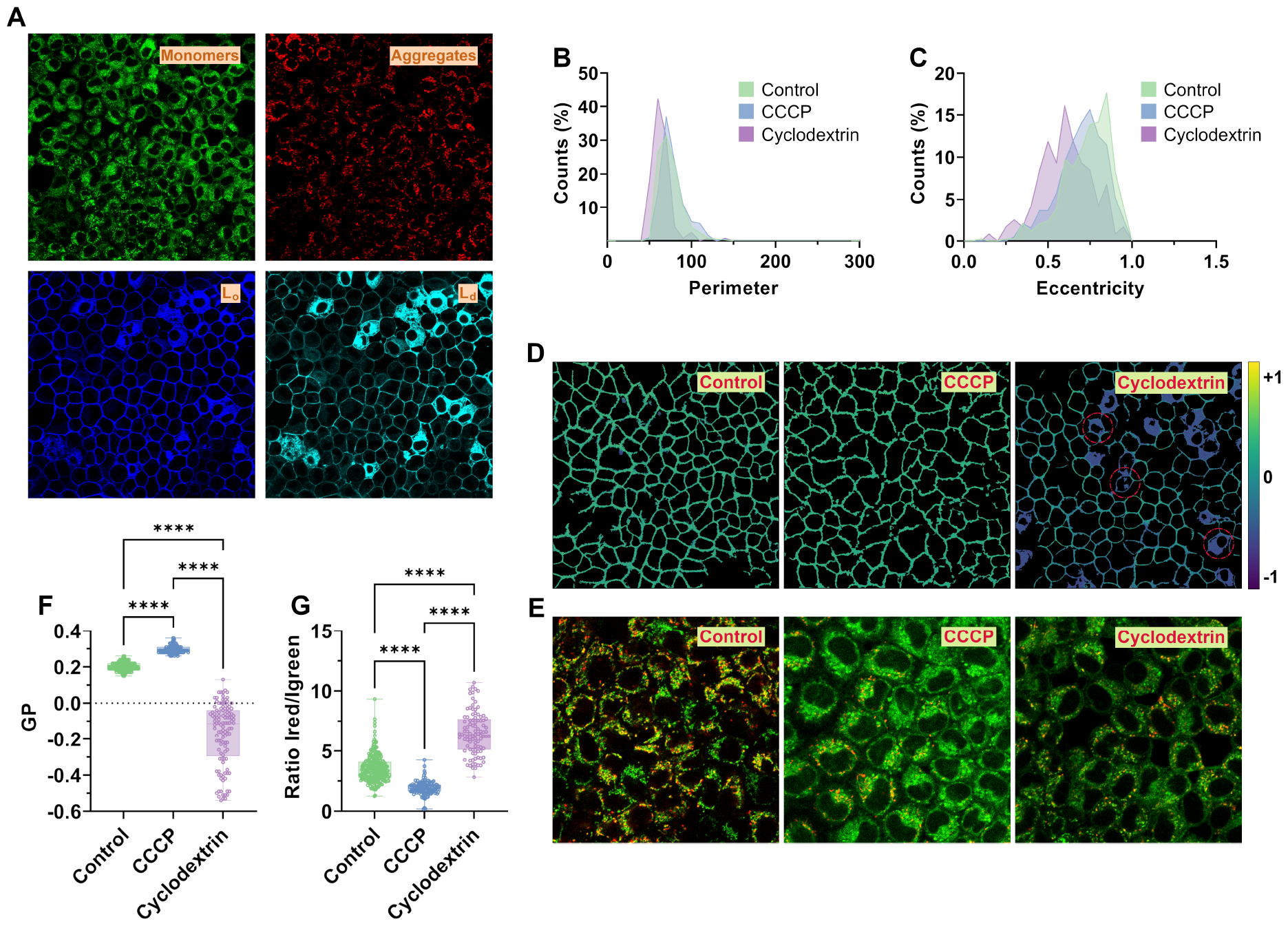
VISION allows for correlation of membrane fluidity with mitochondrial health. (A) Four different intensity channels used to correlate fluidity of the plasma membrane with mitochondria membrane potential. *Lo* (liquid ordered) and *Ld* (liquid disordered) refer to the channels used for the calculation of GP. (B) Distributions of cell perimeters for control and drug-treated samples. (C) Distribution of cell eccentricities for control and drug-treated samples. (D) GP colour-coded images from the three different samples. The colour bar represents the range of GP values. Red circles mark example cells with dye internalization. (E) Images showing the overlap between JC-1 monomers (green) and aggregates (red) from the three different samples. (F-G) Box plots showing the difference in GP (F) and ratio I_red_/I_green_ (*i*.*e*., mitochondrial potential, G) between the different conditions. Statistical differences between samples were assessed *via* ANOVA analysis using the non-parametric Kruskal-Wallis test.

Instead, the blue and cyan channels were used to detect signal from ordered (*L*_*o*_) and disordered (*L*_*d*_) phases, respectively and calculate the GP. Channel range was optimized to avoid spectral spillover of JC-1 into Pro12A channels (Fig. S3). By performing single-cell characterization, we observed changes in cell morphology due to drug treatment. Specifically, both CCCP and Cyclodextrin induced a slight decrease in cell perimeter (Fig. 5B) and eccentricity (*i*.*e*., more rounded cells, Fig. 5C), which could be ascribed to a partial cell detachment from the well’s bottom. Significant changes between control experiments and treated cells were also observed in membrane fluidity and mitochondrial potential (Fig. 5D-E). While CCCP caused a slight decrease in membrane fluidity (*i*.*e*., higher GP values), the removal of cholesterol *via* cyclodextrin drastically lowered the median GP values, in agreement with data in literature (Fernández-Pérez *et al*., 2018). Treatment with the cyclodextrin also promoted dye internalization (red circles in Fig. 5D) consequently to the disruption of lipid bilayer packing. Interestingly, the two drugs had an opposite effect also on the mitochondria membrane potential. CCCP strongly impaired mitochondria health, as shown from a lower ratio I_red_/I_green_ from masked mitochondria and an overall decrease in JC-1 aggregates, as reported in literature (Miyazono *et al*., 2018). On the other hand, cyclodextrin did not affect the number of JC-1 aggregates-per-cell, but instead, it caused a notable increase in mitochondrial potential. This hyperpolarization of mitochondria could be a direct consequence of the cholesterol depletion from the plasma membrane which, in the short term, would trigger recruitment of cholesterol from other organelles’ membranes to counteract the effect.

### Protein accumulation at the contact site between two cells

While we so far used VISION to study biophysical parameters, one is not limited to these and can study any multi-parametric aspect of images. For this propose, we also tested the software applicability in studying cell-cell contact areas, which are critical for intercellular communication, cell polarization and tissue integrity (Garcia, Nelson and Chavez, 2018). Specifically, we focused on the redistribution of signalling proteins at a reconstituted immune synapse (IS). The IS describes a highly spatio-temporally organized contact between immune cells and their target antigen-presenting cells resulting in the activation of immune signalling (Monks *et al*., 1998; Dustin, 2014). The spatial organization of proteins within the IS resembles a bull’s eye-like structure comprising supramolecular activation clusters (SMACs), which are characterized by focal accumulation or exclusion of certain proteins (Dustin, 2014). The enrichment of the two adhesion receptors CD2 and CD58 at the cell-cell contact is important for IS formation and function (Dustin, 2014; Sanchez-Madrid et al., 1982; Shaw et al., 1986). On the other hand, the exclusion of the tyrosine phosphatase CD45 from the IS ensures proper regulation of the immune signalling process (Dustin, 2014; Chang et al., 2016). In Fig. 6, we show the results obtained from a minimal system resembling the IS which consisted in GUVs decorated with CD2 and CD45 (1:1 ratio) in contact with Jurkat T cells. As expected, we observed the enrichment of CD2 and a corresponding exclusion of CD45 at the cell-GUV contact, thus confirming previous observations on similar systems (Céspedes et al., 2021; Jenkins et al., 2018). Interestingly, from the analysis of the two intensity trajectories—green for CD2 and magenta for CD45—we observed a quite broad heterogeneity in protein accumulations and/or exclusion (*i*.*e*., signal intensity) from the contact area and the tendency of CD2 to form focal clusters within the same IS (Figure 6A-C). Therefore, our software can be used to profile cell-to-cell contact sites at a sub-micrometre spatial resolution and study the biological significance of protein clustering/redistribution within those areas in various scenarios.

**Fig. 6.**
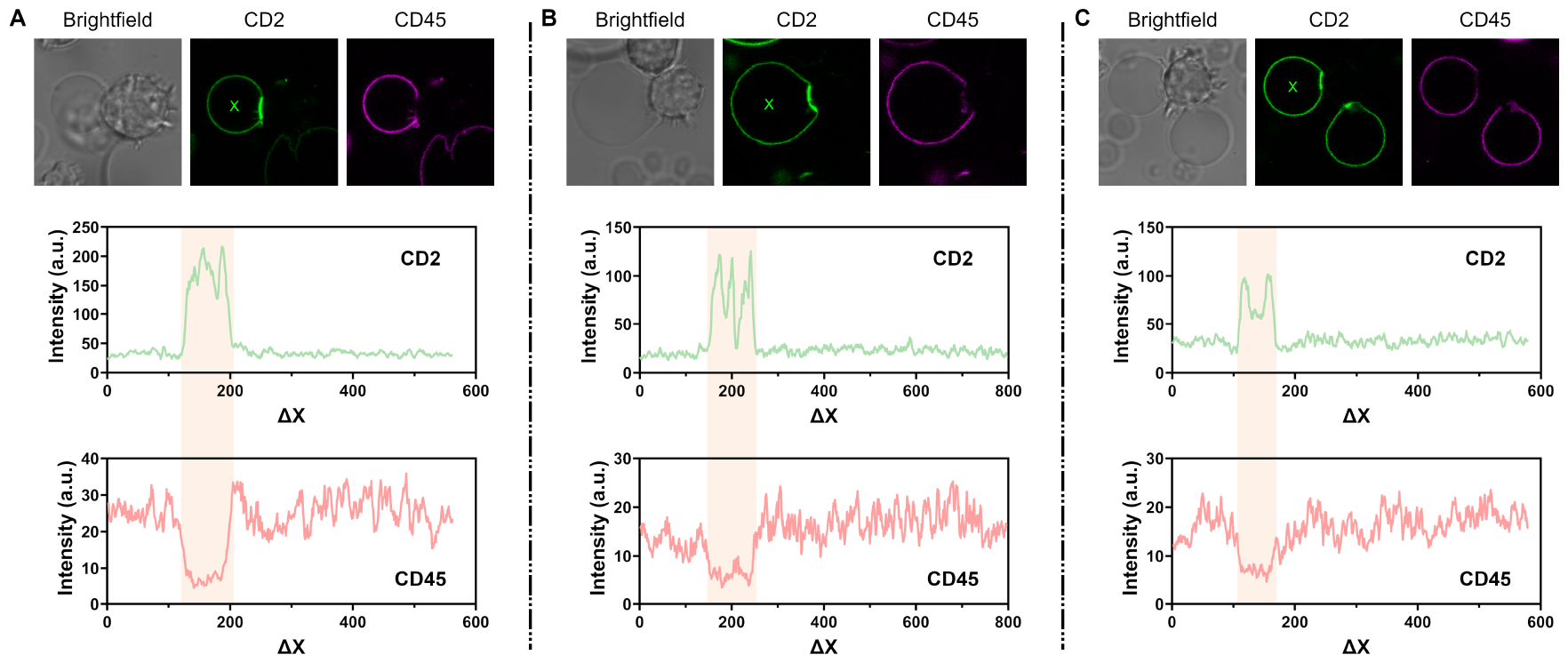
VISION reveals protein cluster formations at the cell-to-GUV contact site. (A-C) Examples of Jurkat T cells interacting with GUVs decorated with CD2 and CD45 adhesion proteins. The green and magenta/red channels refer to CD2 and CD45 protein, respectively. The pink areas in the profiled trajectories highlight either accumulation or exclusion of proteins from the membrane-membrane contact area.

### Profiling nanodomains with super-resolution

As mentioned above (Fig. 3) VISION adopts a different approach to profile the lipid bilayer of vesicles by sliding customizable moving average filter along a membrane-centred guideline. By doing so, it ensures a constant area of integration which is independent from the local membrane morphology. This is crucial to obtain quantitative measurements of small variations in the membrane biophysical properties under investigation. Furthermore, since the size of the integrating element ultimately depends on the image pixel size (with the smallest element being a 3x1 pixels line), by operating at high spatial resolution it is possible to capture heterogeneities occurring at the nanoscale. This is shown in Fig. 7, where we compare the GP profiles measured with either confocal or stimulated emission depletion (STED) microscopy from the same phase separated GPMV. In both scenarios, the pixel size was set to ∼32 nm and a 3x3 pixel averaging filter was chosen for GP integration, thus yielding a total area of ∼0.08 nm^2^ per profiled point (*i*.*e*., spatial displacement ΔX). From the two areas highlighted in Fig. 7B-D, it appears clear how both the spatial resolution and contrast in GP magnitude (*i*.*e*., ΔGP) between phases is enhanced in STED *versus* confocal. Thus, our software can be used to investigate the dynamic of biophysical remodelling of membranes at both the μm- and nm-scale.

**Fig. 7.**
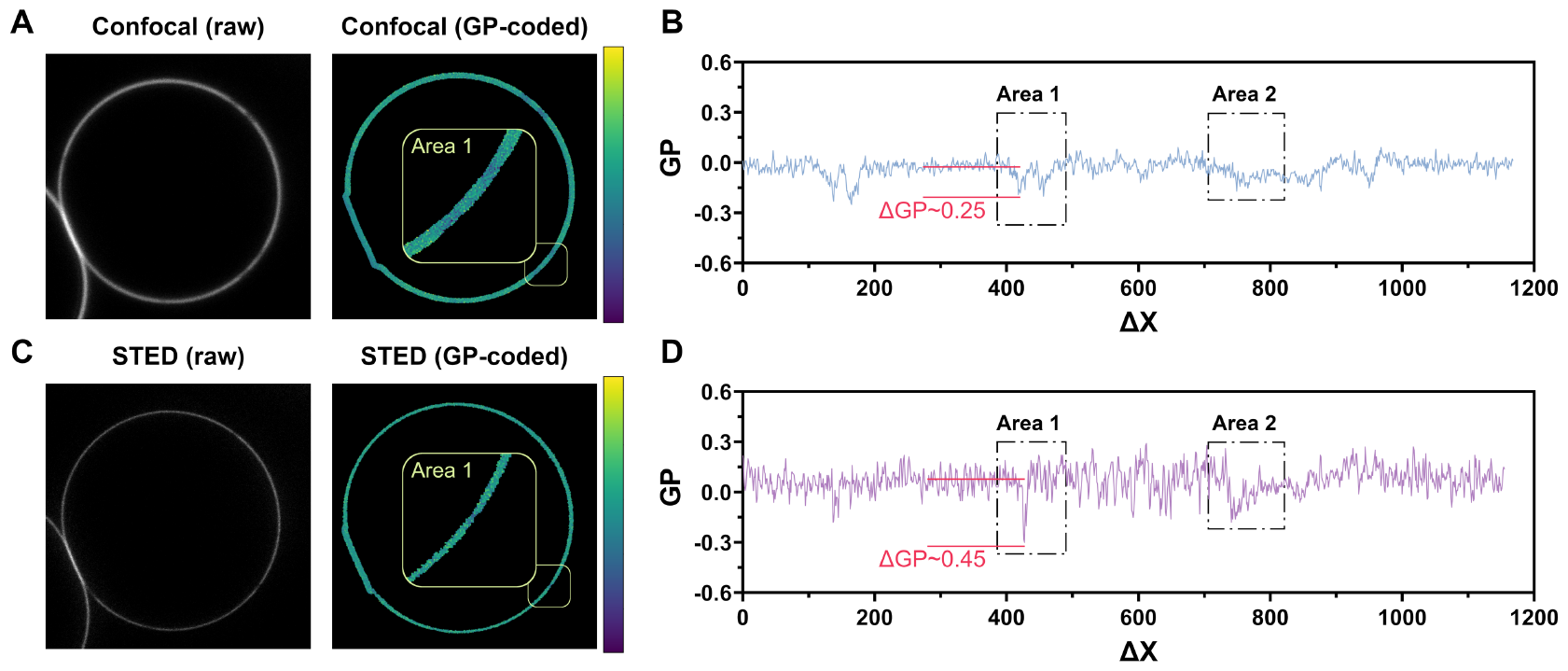
VISION can profile membrane fluidity with super resolution. (A-B) An example of phase-separated GUV stained with Di-4-AN(F)EPPTEA imaged with confocal microscopy (A) and its corresponding profiled trajectory (B). (C-D) An example of phase-separated GUV imaged with STED microscopy (C) and its corresponding profiled trajectory (D). Images reused from *(Sezgin* et al., *2017)*.

## Discussion

With the advancement in technologies, microscopy is quickly overcoming the historical limitation in throughput and dimensionality. New instrumentation offers the possibility to extend the collectable information beyond the spatial and temporal space to the spectral dimension, thus delivering improved multiplexing (Huang *et al*., 2022). VISION was developed to take advantage of the multi-channels and spectral modalities implemented in many commercial microscopes and perform complex mathematical operations from up to four different channels. Thanks to the different modules implemented, our software finds large applicability in a plethora of scenarios which span from correlative analysis between the plasma membrane and the cytosol or subcellular compartments, experiments on proteins colocalization and measurement of cellular properties which require the use of ratiometric environment-sensitive probes, such as those for membrane fluidity, mitochondria potential *etc*. Furthermore, the high versatility of the module for image masking allows to work with a variety of biological samples panning from vesicles (*e*.*g*., GUVs, GPMVs *etc*.) to live suspension and adherent cells. In addition, VISION’s novel approach to membrane profiling enhances the accuracy of signal integration, potentially improving quantitative analysis of, for example, protein clustering/accumulation at the cell-to-cell junctions (Shashikanth, Kisting and Leckband, 2016), spatial/temporal remodelling of membrane biophysical properties and dynamics of phase separation.

### Future developments

Given the increasing interest in studying biophysical properties of cell and organelle membranes, we will extend the capability of our software by introducing extra functionalities to perform adaptive thresholding—which results useful when working with z-stack and t-stack of images with heterogenous brightness—and AI-based segmentation. Currently, segmentation of the image occurs at the object level, to identify individual cells from clusters. In the future, we plan to implement an additional module for intra-cytosol particles detection, which will further broaden the applicability of our software to studies on subcellular structures. Finally, we will introduce the possibility to perform single-particle/single-cell tracking analysis of temporally resolved images, and to perform spectral deconvolution of the emission spectrum, thus increasing the multiplexing capability.

## Materials and Methods

### Licenses, GUI development & standalone executables

For the GUI development and distribution of standalone executables, the software VISION is licensed under GPL-3.0. The standalone software application VISION was developed, featuring a graphical user interface (GUI) implemented using Python and the Qt framework (PyQt). The GUI was designed to facilitate user interaction and streamline analysis tasks, including image loading, preprocessing, analysis algorithms, and result visualization. The software was packaged using PyInstaller to create standalone executables for Windows and macOS platforms, enabling easy deployment and usage across various operating systems. This approach ensures accessibility and usability for researchers in the field of microscopy image analysis, offering a user-friendly solution for their analytical needs.

### Preparation of synthetic vesicles

To generate synthetic vesicles different lipids were mixed at different ratio, namely POPC (Avanti Polar Lipids, cat# 850457C), DOPC (Avanti Polar Lipids, cat# 850375C), Egg Sphyngomyelin (SM, Avanti Polar Lipids, cat# 860061C), Cholesterol (Avanti Polar Lipids, cat# 700000P) and DGS-NTA(Ni) (18:1, Avanti Polar Lipids, cat# 790404C). For normal GUVs, POPC was mixed with 2% of DGS-NTA(Ni), whereas phase-separated GUVs were obtained from the lipid mixture SM:DOPC:Cholesterol 2:2:1. Then, vesicles were generated by electroformation using a custom-built GUV Teflon chambers with two platinum electrodes (Sezgin, Levental, *et al*., 2012) according to a previous protocol (Sezgin, Kaiser, *et al*., 2012). A volume of 6 μL of 1 mg/mL lipid in chloroform was homogeneously distributed on the electrodes and dried under a nitrogen stream. After placing the electrodes in 370 μL of 300 nM sucrose solution (370 μL), electroformation was performed at 2 V and 10 Hz for 1 h followed by 2 V and 2 Hz for 30 min. Phase-separated GUVs were prepared above the specific lipid transition temperature at 70°C, whereas the other GUVs were generated at RT. GPMVs were generated from mouse fibroblast cell line NIH/3T3 following a slightly modified version of a protocol previously described (Sezgin, Kaiser, *et al*., 2012) In brief, cells were washed twice with HBSS (FisherScientific, cat# 15266355) and incubated for 3 hours at 37 °C and 5% CO_2_ with GPMV-vesiculation buffer consisting of 25 mM methanol-free formaldehyde (FisherScientific, cat# 11586711) and 2 mM dithiothreitol (Biorad, cat# 1610610) dissolved in HBSS. After formation, one batch of GPMVs was incubated with 0.6 mM of Deoxycholic Acid (Sigma-Aldrich, cat# D2510) as described previously (Zhou *et al*., 2013) to stabilize phase separation at RT.

### Cells maintenance

Commercial cell lines were purchased from ATTC: HL-60 (cat # CCL-240), NRK-52E (cat# CRL-1571), Jurkat (clone E6-1, cat# TIB-152), NIH/3T3 (cat# CRL-1658).

Cells were cultured in either RPMI with L-glutamine (FisherScientific, cat# 12004997)—for suspension cells—or DMEM with high glucose (ThermoFisher, cat# 11965092)—for adherent cells—supplemented with 10% Fetal Bovine Serum (Sigma-Aldrich, cat# F7524) at 37 °C and 5% CO_2_. Before experiments on cell-to-GUV contacts, cells were washed twice in PBS at 1500 rpm for 1 min at RT and subsequently counted. For imaging, ∼3 x10^4^ cells per well were added to each well containing GUVs in μ-Slides (18-well glass bottom, ibidi, cat# 81817). For GPMVs preparation, cells were seeded into 12-well plates to reach a confluency of 70-80% on the day of experiment and transferred into an 8-well chambered cover glass (Cellvis, cat# C8-1.5H-N) precoated with Fetal Bovine Serum Albumin (Sigma-Aldrich, cat# A3803). For drug reatment experiments, NRK cells were seeded into 8-well polymer bottom Ibidi slide at 70% confluency (#1.5, cat# 80801) a day before the analysis.

### Protein labelling of GUVs

The proteins CD2 (Human, Recombinant, ECD, His Tag from SinoBiological, cat# 10982-H08H) and CD45 (Human, Recombinant, ECD, His Tag from SinoBiological, cat# 16884-H08H) were labelled with Alexa Fluor™ 488 NHS Ester (Succinimidyl Ester, ThermoFisher, cat# A20000) and Alexa Fluor™ 647 NHS Ester (Succinimidyl Ester, ThermoFisher, cat# A37573), respectively. The amount of NHS Ester dye needed for the labelling reaction was calculated following Eq.1:

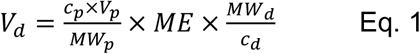

where *V*_*d*_ is the volume of the dye solution, *c*_*p*_ is the protein concentration, *V*_*p*_ is the volume of protein, *MW*_*p*_ is the protein molecular weight, *ME* is the molar excess (number of dye molecules per protein molecule, here we chose 5) and *c*_*d*_ is the concentration of the dye solution. To each 1 mL protein solution, 0.1 mL NaHCO_3_ (1 M) solution in water pH 8.5 was added. The calculated amount of dye solution (in DMSO) was added to the protein solution and incubated at RT for 1 hour with continuous stirring. The protein-dye solution was then added to two consecutive 0.5 mL 40K MWCO Zeba™ Spin Desalting Columns (ThermoFisher, cat# 87767) (previously washed twice with PBS) and centrifuged at 1500xg for 2 min.

### Drug treatments

For experiments on cholesterol depletion, a 10 mM solution was prepared by dissolving 0.12 g of methyl-beta-cyclodextrin in 10 mL L-15 medium (Sezgin *et al*., 2015). NRK-52E cells were then incubated for 30 minutes at 37 °C and 5% CO_2_ in 300 μL/well of cyclodextrin solution together with JC-1. For experiments on disruption of the mitochondrial membrane potential, CCCP was added to cells at a final concentration of 20 μM and incubated at 37 °C and 5% CO_2_ for the last 5 minutes of incubation with JC-1 (Sivandzade, Bhalerao and Cucullo, 2019). After treatments, cells were washed twice with PBS, and 300 μL/well of fresh L-15 was added before imaging.

### Labelling of membranes and mitochondria

Phase-separated GUVs were stained at a final concentration of 100 nM of NR12S or NR12A and 300 nM Pro12A, whereas phase-separated GPMVs were labelled with 400 nM of Pro12A. Images were acquired after ∼5 min from staining. For confocal imaging of cell-to-GUV contacts, the GUVs were stained with 0.05 μg of both CD2-488 and CD45-AF647 at RT for 30 min. NRK-52E cells were stained with JC-1 at a final concentration of 800 nM for 3x10^5^ cells/well in Leibovitz’s L-15 medium (ThermoFisher, cat# 21083027) and incubated for 30 minutes at 37 °C and 5% CO_2_ (Sivandzade, Bhalerao and Cucullo, 2019). Then, cells were washed twice with fresh L-15 buffer and stained with NR12A at a final concentration of 500 nM.

### Confocal and STED imaging

Confocal images were acquired using a Zeiss LSM 780 confocal microscope with a 32-channel array of gallium arsenide phosphide (GaAsP) detectors or a Zeiss LSM 980 confocal microscope AIRY2/FCS (for imaging of cell-to-GUV contact areas). For the LSM 780, the wavelengths range for each detector was set to 9 nm. Pro12A was excited using a 405 nm laser, whereas NR12A and NR12S were excited using a 488 nm laser. Different ranges of wavelengths were acquired for the different membrane-intercalating probes, namely ∼423– 601 nm for Pro12A and ∼503–700 nm for NR12A and NR12S (Ragaller *et al*., 2022). For experiments on cell-to-GUV contacts, CD2-AF488 was excited at 488 nm CD45-AF647 at 639 nm. For experiments on mitochondria potential, JC-1 dye was excited with a 488 nm laser and the detection range was set either between 500–550 nm (for JC-1 monomers) or 580– 610 (for JC-1 aggregates). Pro12A was excited with a 405 nm laser and the detection range was set either between 420–510 nm (for Lo) or 470–510 (for Ld).

STED images were acquired using a Leica SP8 3X STED microscope and exciting the environment-sensitive dye Di-4-AN(F)EPPTEA with 488 nm laser and acquiring the emission in two different channels with wavelengths range 520–570nm and 620–700-nm. 775 nm STED laser (∼200 mW measured at the back aperture of the objective) was applied.

### Statistical analysis

Statistical analysis on phase-separated GPMVs and GUVs was done in Prism GraphPhad Version 9.4.1.

## Acknowledgements

We thank the SciLifeLab Advanced Light Microscopy facility and National Microscopy Infrastructure (VR-RFI 2016-00968) for their support on imaging.

## Author contributions

L.A. and F.W. conceived and developed the software. L.A. wrote the core Python algorithms used by VISION, while F.W. developed the Graphical User Interface (GUI) and standalone executables builds. S.I. performed the experiments on correlation between membrane fluidity and mitochondria potential. J.S. performed the experiments on phase-separated GPMVs. F.R. performed the experiments on GUVs and cell-to-GUVs contact. E.S. performed the experiments on confocal *vs* super-resolution microscopy. B.P. and E.S. obtained funding and performed supervision. L.A wrote the original draft, and all authors revised the manuscript.

## Data availability

All data in this manuscript will be publicly available at FigShare repository upon publication. VISION software and the python core-code can be freely downloaded from the GitHub repository (https://github.com/biosciflo/VISION).

